# Generalized paradoxical effects in excitatory/inhibitory networks

**DOI:** 10.1101/2020.10.13.336727

**Authors:** Kenneth D. Miller, Agostina Palmigiano

## Abstract

An inhibition-stabilized network (ISN) is a network of excitatory and inhibitory cells at a stable fixed point of firing rates for a given input, for which the excitatory subnetwork would be unstable if inhibitory rates were frozen at their fixed point values. It has been shown that in a low-dimensional model (one unit per neuronal subtype) of an ISN with a single excitatory and single inhibitory cell type, the inhibitory unit shows a “paradoxical” response, lowering (raising) its steady-state firing rate in response to addition to it of excitatory (inhibitory) input. This has been generalized to an ISN with multiple inhibitory cell types: if input is given only to inhibitory cells, the steady-state inhibition received by excitatory cells changes paradoxically, that is, it decreases (increases) if the steady-state excitatory firing rates decrease (increase).

We generalize these analyses of paradoxical effects to low-dimensional networks with multiple cell types of both excitatory and inhibitory neurons. The analysis depends on the connectivity matrix of the network linearized about a given fixed point, and its eigenvectors or “modes”. We show the following: (1) A given cell type shows a paradoxical change in steady-state rate in response to input it receives, if and only if the network with that cell type omitted has an odd number of unstable modes. Excitatory neurons can show paradoxical responses when there are two or more inhibitory subtypes. (2) More generally, if the cell types are divided into two nonoverlapping subsets A and B, then subset B has an odd (even) number of modes that show paradoxical response if and only if subset A has an odd (even) number of unstable modes. (3) The net steady-state inhibition received by any unstable mode of the excitatory subnetwork will change paradoxically, *i.e.* in the same direction as the change in amplitude of that mode. In particular, this means that a sufficient condition to determine that a network is an ISN is if, in response to an input only to inhibitory cells, the firing rates of and inhibition received by all excitatory cell types all change in the same direction. This in turn will be true if all E cells and all inhibitory cell types that connect to E cells change their firing rates in the same direction.

## Introduction

An inhibition-stabilized network (ISN) is a network of excitatory (E) and inhibitory (I) neurons at a stable fixed point or steady state of firing rates for a given input, in which the E population would be unstable if I firing rates were frozen at their fixed point values, but the fixed point is stabilized by dynamic I responses (*i.e.*, by feedback inhibition) (Ozeki et al., 2009). Tsodyks et al. (1997) showed that, given a model of single E and I populations, ISNs exhibit a “paradoxical” effect: if external excitatory input is added to the I population, the I firing rates paradoxically are decreased in the new steady state.

To review the basis for this, we consider a nonlinear rate equation 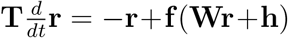 where 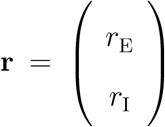 is the vector of E (*r*_E_) and I (*r*_I_) firing rates, 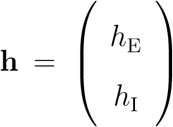 is the vector of external inputs to the network, and **W** is the two-by-two weight matrix, 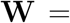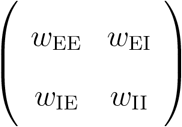, where *w*_XY_ represents the connection strength from population *Y* to population *X*, *X, Y* ∈ {*E, I*}, with *w*_XE_ > 0 and *w*_XI_ < 0. Here, 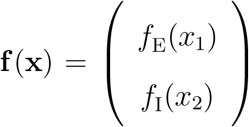 for two monotonically increasing functions *f*_E_ and *f*_I_, and 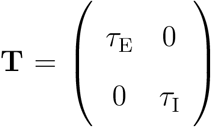 is the diagonal matrix of E and I time constants.

We assume the system has a stable fixed point, and linearize about the fixed point. We let *δ***r** and *δ***h** be the deviations of firing and external inputs from the fixed point values, and let 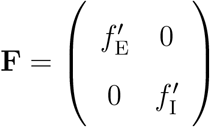 where 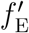 and 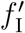 are the derivatives of *f*_E_ and *f*_I_ evaluated at the fixed point. The equation linearized about the fixed point is then

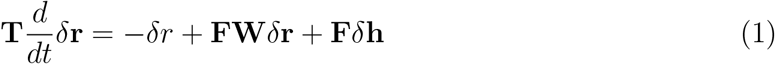

This equation has the fixed point solution

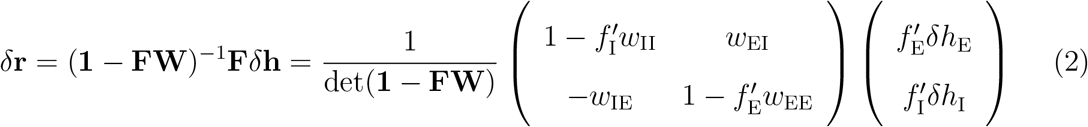

where **1** is the identity matrix (we will use **1** for the identity matrix of any dimension, where the dimension is inferred from the context; here, it is two-dimensional). Inhibition shows a paradoxical response when the sign of *δr*_I_ is opposite to that of *δh*_I_ – *e.g.*, a positive input to inhibitory cells causes a negative steady-state response in those cells. The fixed point is stable, so both eigenvalues of the Jacobian **T**^−1^(**FW**−**1**) have real parts < 0, which implies that det **T**^−1^(**1**− **FW**) > 0 and thus that det(**1**− **FW**) > 0. Thus, given that 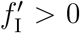, the inhibitory response is paradoxical precisely when 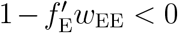. This condition, 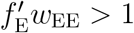, is precisely the condition that the excitatory subnetwork be unstable by itself; that is, that if inhibitory firing is frozen at the fixed-point level (*δr*_I_ = 0), the linearized equation for *r*_E_ exponentially moves away from the fixed point after any perturbation of *r*_E_ from its fixed point value. Thus, inhibition shows a paradoxical response precisely when the excitatory subnetwork is unstable by itself. Evidence has been found that various cortical areas are ISNs (Adesnik, 2017; Kato et al., 2017; Li et al., 2019; Ozeki et al., 2009; Sanzeni et al., 2020).

Recently much attention has focused on the existence of multiple subtypes of I interneurons and their specific connectivity (*e.g.*, Pfeffer et al., 2013). Rubin et al. (2015, Section S2.2.3) and Litwin-Kumar et al. (2016) both showed how the ISN paradoxical result generalized to this case, with a single E unit and multiple I units. The generalization is that, in response to an input only to inhibitory cells, the summed inhibition received by the E unit must change paradoxically, increasing (decreasing) if the added input causes *r*_E_ to increase (decrease). Note that this does not determine the change in the firing rate of any individual inhibitory population.

Rubin et al. (2015) gave a conceptual argument for this. The E subnetwork being unstable by itself means that a perturbation of *r*_E_ above (below) its fixed-point firing rate would recruit too much (lose too much) recurrent excitation, pushing *r*_E_ even higher (lower) in the absence of changes in other inputs to the E unit. Therefore, if external input is added to the network causing *r*_E_ to be increased (decreased) at the new steady state, the input to E other than the recurrent excitatory input must change in a direction that would push *r*_E_ to decrease (increase), to cancel the excess change in recurrent excitation and thus stabilize the perturbed firing rate. In particular, if external input is added only to I cells, then the change in inhibition received by the E unit must be paradoxical.

Litwin-Kumar et al. (2016) independently gave a mathematical version of this argument. Starting with the linearized equation for the steady state response to the perturbation *δ***h**, (**1**− **FW**)*δ***r**= **F***δ***h**, and given that the external input that perturbs from the fixed point is given only to the I populations (*δh*_E_ = 0), the equation for *r*_E_ reads

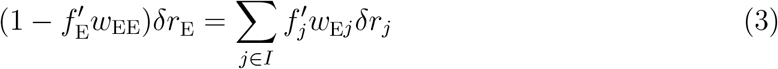

where *j* ∈ *I* indicates that *j* is one of the I populations, and *δr*_*j*_ is the perturbation from the fixed point of the firing rate of this I population, which connects to the E population with strength *w*_E_*j* < 0 and has nonlinearity *f*_*j*_. The right hand side of Eq. 3 is the total change in inhibitory input to the E cells (relative to that received at the fixed point): if it is negative, the inhibition received has increased. The left side has the same sign as *δr*_E_ in a non-ISN, but the opposite sign as *δr*_E_ in an ISN. Thus, in an ISN, if *δr*_E_ is negative (positive), there is a paradoxical net decrease (increase) in the inhibition received by the E population. Note that, in the expression 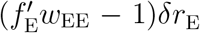, the −*δr*_E_ term represents the change in E leak current induced by *δr*_E_, while the 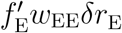 term represents the change in recurrent excitation to E cells induced by *δr*_E_. Thus, if 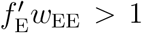, it means the perturbation *δr*_E_ induces too large a change in recurrent excitatory input – more than is needed to compensate the change in leak current – causing *δr*_E_ to grow even larger in magnitude in the absence of inhibitory response; and this excess change in recurrent excitatory input must be cancelled by a paradoxical change in the inhibition received to stabilize the change in firing rate *δr*_E_. This is the mathematical version of the conceptual argument in Rubin et al. (2015).

In this paper, we generalize these results to arbitrary numbers of *E* and *I* celltypes. We determine when a paradoxical firing-rate response of a given cell type or pattern across cell types will arise. We state the generalization of Eq. 3: what constitutes a paradoxical change in input received when there are multiple *E* and *I* subtypes? At some points we will refer to the much-studied 4-cell-type case of E cells and three types of I cells: parvalbumin-expressing (*P*), somatostatin-expressing (*S*), and VIP-expressing (*V*) (Pfeffer et al., 2013). Here, we use one unit for each cell type. In Palmigiano et al. (2020), we address the effects of having many, inhomogeneous neurons of each cell type.

## Results

We represent each cell type by one unit. We consider a network with *N* = *N*_E_ + *N*_I_ units: *N*_E_ E units and *N*_I_ I units. We assume that the network is at a stable fixed point, and we consider the linearized dynamics about the fixed point. For simplicity, we write **r** instead of *δ***r** and **h** instead of *δ***h**, so the linearized dynamics are

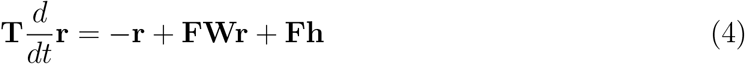

Here, 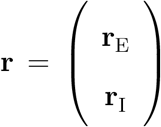, where **r**_E_ and **r**_I_ are the *N*_E_-dimensional and *N*_I_-dimensional column vectors of E-cell and I-cell firing rates respectively. By unit *j*, we mean the unit whose firing rate is the *j*^*th*^ element of **r**. The element of **W** representing the connection from unit *k* to unit *j* is written *w*_*jk*_, where again *w*_*jk*_ > 0 for *k* an excitatory unit and *w*_*jk*_ < 0 for *k* an inhibitory unit.

We will also consider alternative coordinate systems in which elements of **r** may be the firing rate of various linear combinations or patterns of the elements of **r** that do not mix the E and I units, and **W** and **h** are correspondingly transformed to represent effective connections between, and inputs to, these patterns. In particular, we may consider using the eigenvectors of the E submatrix instead of the E units, and/or use the eigenvectors of the I submatrix instead of the I units. We will often use “modes” to mean eigenvectors and will use the two terms interchangeably. We will more generally refer to “units”; in most cases (except where a particular coordinate system is specified), the results apply in any coordinate system and so could refer to patterns rather than the units representing individual cell types.

### 1. Criterion for a single unit to show a paradoxical response

The steady-state response to the perturbation **h** is given by (**1**− **FW**)^−1^**Fh**, that is, (**1**− **FW**)^−1^**F** determines the response to a given perturbation **h**. Hence we call **R**≡ (**1**− **FW**)^−1^**F** the response matrix; this can also be written (**F**^−1^−**W**)^−1^. Unit *i* has a paradoxical firing-rate response if the corresponding diagonal element of the response matrix, *R*_*ii*_, is negative.

The elements of the matrix inverse are given by 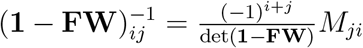 where *M*_*ji*_ is the (*j, i*) minor of **1**− **FW**, meaning the determinant of the submatrix formed by deleting the *j*^*th*^ row and *i*^*th*^ column of **1**− **FW**. Thus, since **F** is diagonal, the diagonal entries of *R* are

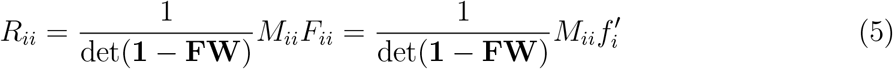

Given that 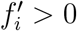, then if det(**1**− **FW**) > 0, the response of unit *i* is paradoxical iff *M*_*ii*_ < 0.

As noted in Palmigiano et al. (2020), this is closely related to the stability of the network with unit *i* deleted. The Jacobian, the matrix multiplying **r** in the linearized equation for 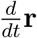, is **J** = **T**^−1^(**FW** – 1) = −**T**^−1^**FR**^−1^, so **R**= (−**J**)^−1^**T**^−1^**F**. Stability of the fixed point is equivalent to all of the eigenvalues of **J** having negative real parts, or all the eigenvalues of −**J** having positive real parts. Therefore, since we are assuming the fixed point is stable, det(−**J**) > 0. But det(−**J**) = det(**T**^−1^) det(**1**− **FW**); since **T**^−1^ is a diagonal matrix with positive elements, its determinant is positive, so from det(−**J**) > 0 we conclude det(**1**− **FW**) > 0. Thus, *R*_*ii*_ < 0 iff *M*_*ii*_ < 0. By the same formula as above for the inverse of a matrix, the diagonal elements of (−**J**)^−1^ are given by 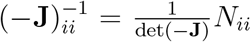 where *N*_*ii*_ is the (*i, i*) minor of −**J**. *N*_*ii*_ is the determinant of the negative of the Jacobian of the network with unit *i* omitted. Thus, if the network with unit *i* omitted is stable, then *N*_*ii*_ > 0. Finally, we note

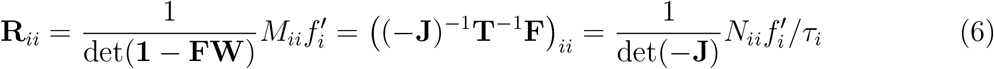

All of the factors 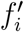, *τ*_*i*_, det(−**J**), and det(**1**− **FW**) are positive. Thus, *R*_*ii*_ < 0 ↔ *M*_*ii*_ < 0 ↔ *N*_*ii*_ < 0, and *N*_*ii*_ > 0 if the subcircuit with unit *i* omitted is stable.

More generally, *N*_*ii*_ > 0 iff the sub-circuit with unit *i* removed has an even number of unstable modes (which includes the case of 0 unstable modes), and *N*_*ii*_ < 0 iff that subcircuit has an odd number of unstable modes. Thus, we arrive at our first result: **The***i*^*th*^ **unit in a network has a paradoxical response to its own stimulation if and only if the subnetwork with the***i*^*th*^ **unit excluded – the dynamics with***r*_i_ **held fixed at its fixed point value – has an odd number of unstable modes.** This in turn is true iff the minor *M*_*ii*_ of the matrix **1**− **FW** is negative, that is, iff the matrix **1**− **FW** with the *i*^*th*^ unit omitted has an odd number of unstable modes, meaning modes with negative real eigenvalue.^1^

In the two-dimensional case – one E unit, one I unit – the minor of **1**− **FW** for the *E* unit is 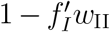, which is always positive, so the E unit cannot have a paradoxical response. The minor of the *I* unit is 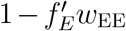, so the I unit has a paradoxical response precisely when 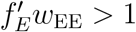, *i.e.* when the E unit is unstable by itself.

More generally, with 1 E unit, the minor of the E unit is just the determinant of the I submatrix of **1**− **FW**. Thus, with more than one I unit, as in the 3-I-subtype models mentioned above, the E subunit can show a paradoxical response: although all the elements of the I submatrix of **1**− **FW** are positive, the submatrix can have negative eigenvalues if it is more than 1-dimensional. Such a paradoxical E response was first shown in (though not noted in) Garcia Del Molino et al., 2017, Fig. 2d. An example of a network with a paradoxical E response is shown in Fig. 1.

**Figure 1:**
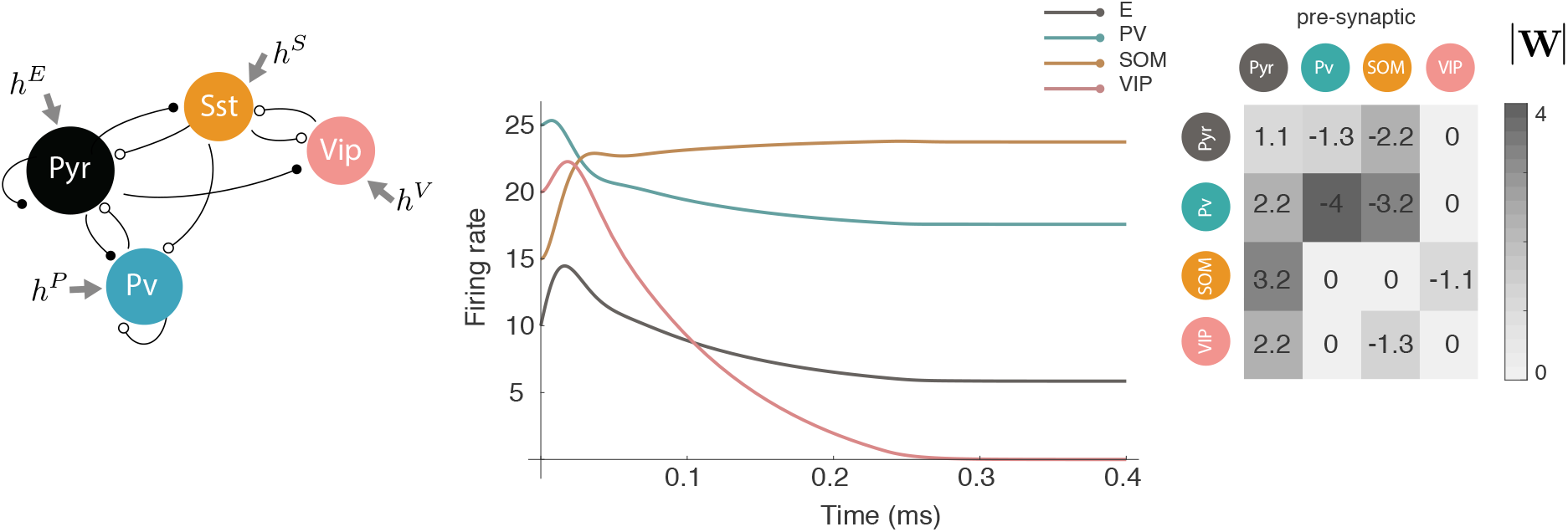
Paradoxical excitatory response. In a model with one excitatory (*E*) unit and three inhibitory units, labelled *P*, *S* and *V*, an input to *E* given at *t* = 0 causes a paradoxical decrease in *E* firing rate in the new steady state. Neurons were given inputs that made the firing rates shown at *t* = 0 a steady state; then at *t* = 0 an additional excitatory input was added to the E unit. Dynamics were linear rectified: 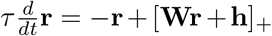 where [**x**]_+_ means the vector **x** with negative elements set to zero, and *τ* = 20ms. The connectivity matrix was 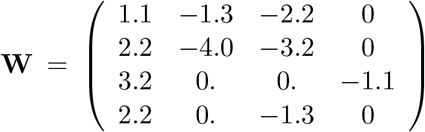 where rows or columns 1 through 4 indicate *E*, *P*, *S*, *V* respectively, with rows representing the postsynaptic and columns representing the presynaptic cell. This matrix was designed simply to illustrate a paradoxical *E* response and is not meant to model the actual connectivity among these cell types. Inputs **h** to *E, P, S, V* to yield initial steady state were (64.5, 151., 5., 17.5); then at *t* = 0, the input to *E* was increased by 10.

Furthermore, as pointed out in (Mahrach et al., 2019), with multiple inhibitory neuron types, a paradoxical response in one inhibitory neuron type need not imply that the excitatory subnetwork alone is unstable, *i.e.* that the network is an ISN. Instead, the requirement that, for example, parvalbumin (*P*) neurons give a paradoxical response in the 3-I-subtype model is that the subnetwork composed of the other three types – *E*, somatostatin-expressing inhibitory neurons (*S*), and VIP-expressing inhibitory neurons (*V*) – be unstable, which does not require that *E* alone be unstable. To illustrate, let 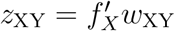, where *X* and *Y* may take the values *E*, *P*, *S*, or *V*. Using the common assumption that *w*_EV_ = *w*_PV_ = *w*_SS_ = *w*_VV_ = 0 (based on the relatively small values of these connections in Karnani et al., 2016; Pfeffer et al., 2013; *w*_SP_ also is small in Pfeffer et al., 2013 but not in Karnani et al., 2016) and the assumption that the fixed point is stable, the *PP* element of the response matrix has the same sign as 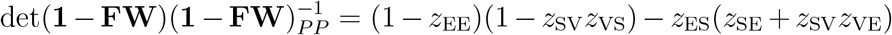. Parvalbumin neurons show a paradoxical response when this element is negative, which does not require excitatory instability (excitatory instability corresponds to (1 − *z*_EE_) < 0). Note that this expression does not involve any connections to or from *P* neurons, since these are not involved in determining whether or not the *E/S/V* circuit is unstable.

An experiment that does not measure the inhibitory currents received by excitatory neurons can nonetheless establish that a network is an ISN if it determines that an input to one set of inhibitory neurons causes *all* inhibitory cell types that connect to *E* cells to change their firing rates in the same direction as *E* cells. This observation is sufficient to establish an ISN because it establishes that the net inhibition received by *E* cells changed paradoxically, *i.e.* in the same direction as the change in *E* cell firing rates. For example, *a paradoxical response of parvalbumin (P) neurons, along with the observation that E cells and all other inhibitory cell types that connect to E cells all change their firing rates in the same direction as the P cells when the P cells are stimulated*, together would suffice to demonstrate that the network is an ISN.

### 2. More general relationship between the number of unstable modes in one set of units and paradoxical responses in the complementary set of units

Suppose we divide the set of units into two complementary sets, *e.g.* the excitatory-cell subtypes and the inhibitory-cell subtypes. Then we can write **1**− **FW** as a block matrix, 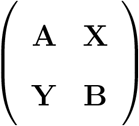, where **A** represents the subnetwork of the first set of units, **B** the subnetwork of the second set of units, and **X** and **Y** represent the connections between the two sets. Then, assuming **A** is invertible, we have the following well-known identity for block matrices:

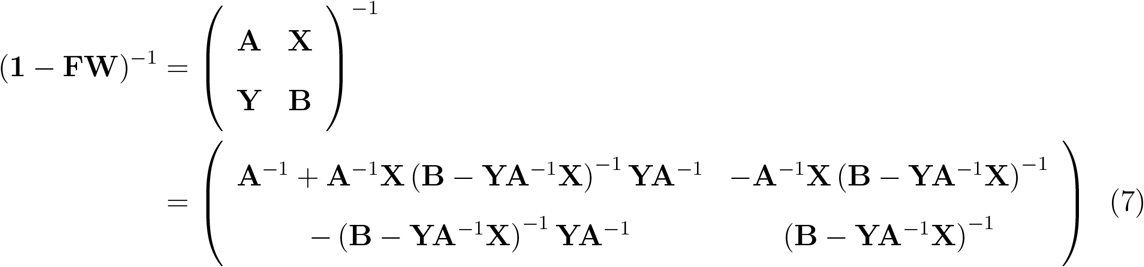

We can now multiply **1**− **FW** by a matrix with unit determinant to obtain a new matrix with the same determinant as **1**− **FW**:

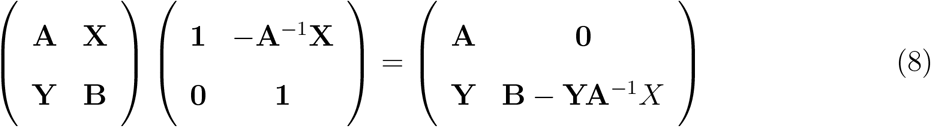

(Here, **0** means a matrix of all zeros, with size determined by the context.) This means that

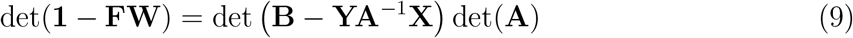

We know that det(**1**− **FW**) > 0, so this tells us that det (**B**− **YA**^−1^**X**) and det(**A**) have the same sign (and det (**B**− **YA**^−1^**X**)^−1^ always has the same sign as det (**B**− **YA**^−1^**X**)). From Eq. 7, we see that (**B**− **YA**^−1^**X**)^−1^ is the response matrix for the subcircuit **B**. Thus, either **A** and (**B**− **YA**^−1^**X**)^−1^ both have an even number of modes with negative real eigenvalues, or they both have an odd number of such modes. Such modes are unstable modes of **A**, and are modes of (**B**− **YA**^−1^**X**)^−1^ that show paradoxical response.

This yields the following relationship: **Suppose we separate a circuit into two complementary subsets of units. Then if the subcircuit corresponding to one subset has an even (odd) number of unstable modes, the submatrix of the response matrix corresponding to the other set of units has an even (odd) number of modes with paradoxical response.**

This generalizes our previous result, in which we divided the matrix into one subset containing the single unit *i* and a complementary subset containing all of the remaining units. There, we found that if the remaining units had an odd (even) set of unstable modes, then unit *i* responded paradoxically (non-paradoxically), *i.e.* unit *i*’s “submatrix” of the response matrix (the single diagonal entry *R*_*ii*_) has 1 (0) modes with paradoxical response.

### 3. Paradoxical changes in received input from the opposite type in response to input to the opposite type

Now suppose we transform the E→E submatrix of (**1**− **FW**) to its eigenvector basis, so that it is a diagonal matrix with the eigenvalues on the diagonal. We leave the basis for the I units unchanged. We refer to the *jk* element of **FW** in this new basis as 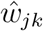. Let us use Greek letters to index units in the eigenvector basis. Suppose a given excitatory mode of (**1**− **FW**), say mode *α* ≤ *N*_E_, has a real^2^ negative eigenvalue *λ*_*α*_ (eigenvalue of the E→E submatrix), meaning the mode is unstable by itself. Then, just as in the argument of Litwin-Kumar et al. (2016) above, if an input is given only to I cells (units *N*_E_ + 1 to *N*), then for the steady state we can write

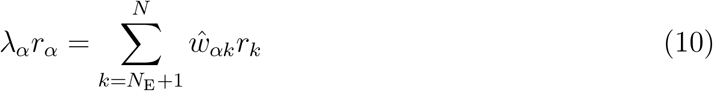

(note that there is no contribution from excitatory cells on the right side because the E→E submatrix is diagonal). Since the eigenvalue *λ*_*α*_ is negative, any change in *r*_*α*_ is accompanied by a paradoxical change of opposite sign in the right-hand side of Eq. 10, which we will see represents the summed input from inhibitory cells to excitatory mode *α*. For example, if *r*_*α*_ decreases, the right side of Eq. 10 would be positive, representing a paradoxical decrease in the inhibition received by excitatory mode *α*.

To see why the right side of Eq. 10 represents the summed input from inhibitory cells to excitatory mode *α*, we define **c** to be the *N*_E_-dimensional vector whose *i*^*th*^ element is the summed inhibition received by the *i*^*th*^ excitatory cell, 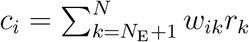. Then, as derived in the Appendix, the right side of Eq. 10 is precisely the component of **c** in the direction of mode *α*, *i.e.* in the basis of E-eigenvectors it is **c**_*α*_, which is what is meant by the summed inhibitory input to mode *α*. This in turn is given by the dot product of the left eigenvector of mode *α* with **c**.

To summarize: **Suppose an eigenvector of the excitatory subnetwork is unstable by itself, with a real negative eigenvalue. Then, if a stimulus is given only to inhibitory cells, the net inhibition to that eigenvector will change paradoxically; that is, the net inhibition received will change in the same direction as the change in the eigenvector’s amplitude.**

In practice, this result is likely to be useful only for excitatory modes that have all elements of the same sign, in the original basis of E- and I-cell types; *i.e.* representing a mode of E cell activity in which all E cell types change their firing rates in the same direction. This is because it is likely to be impractical to measure the change in amplitude of, and change in inhibitory or excitatory input to, a mode in which different elements have different signs (*i.e.*, an activity mode in which some E cell types increase their firing rates while others decrease their rates).

This can be seen as follows. A mode of the excitatory subnetwork of **FW** is unstable if it has an eigenvalue (of that submatrix of **FW**) with eigenvalue > 1 (so that the mode has an eigenvalue of that submatrix of **1**− **FW** that is < 0). The excitatory submatrix of **FW** can be assumed to have only positive entries.^3^ Therefore, by the Perron-Frobenius theorem, there is a single mode with the largest (*i.e.*, most positive) eigenvalue real part, and this eigenvalue is real; and the corresponding right and left eigenvectors, chosen with suitable overall sign, have all entries positive (and they are the only all-positive right and left eigenvectors). If any excitatory modes are unstable, then in particular this all-positive mode will be unstable. This mode represents all excitatory cell types changing their firing rates in the same direction, and the inhibition received by this mode represents inhibitory input of the same sign to all excitatory cell types.

This provides a practical means to determine that the all-positive mode is unstable, meaning that the network is an ISN, without knowing the structure of the all-positive mode. If a perturbation of inhibitory cells leads all excitatory cell types to change their firing rates in the same direction and to all have a change in the inhibition they receive in that same direction, then we can be certain that this all-positive mode and the inhibition it receives both change in the same direction. This is because the change in the mode’s amplitude is the dot product of the all-positive left eigenvector with the single-signed changes in excitatory rates, while the change in the inhibition the mode receives is the dot product of the all-positive left eigenvector with the single-signed changes in inhibition received by the excitatory cells. Both of these quantities will thus change in the same direction. Note that this is a sufficient condition to conclude that the all-positive mode is unstable, not a necessary condition. This is because the two dot products could have the same sign even if the changes in rates of and/or in inhibition received by excitatory cells do not all have the same sign.

We could make a similar argument that, if a mode of the inhibitory submatrix is unstable, and an input is given only to excitatory cells, the excitation received by the unstable inhibitory mode must paradoxically change in a direction opposite to the change in the mode’s amplitude. However, this is not likely to be of practical use. This is because the entries of the I→I submatrix are all negative, and so by the Perron-Frobenious theorem the mode with all entries of the same sign has the eigenvalue with largest *negative* real part, and so it is necessarily stable. So any unstable modes of the inhibitory submatrix represents different inhibitory units changing their activities in different directions. Without knowledge of the structure of this mode (and its left eigenvector), it would be in practice impossible to determine if the excitation it receives changes paradoxically.

More generally, given any subset of units, if a mode of that subset is unstable, and an input is given only to the complementary subset of units, then the input the mode receives from the complementary subset must change in a direction that would drive the mode’s amplitude in the opposite direction to its actual change. This is necessary to compensate for the too-large change in the mode’s input onto itself, which would drive it further in the same direction as its actual change. Again, it is difficult to see how this could be of practical use, except in the special case in which the subset consists of excitatory cells.

In sum, the results of this section give a sufficient, though not necessary, condition for concluding that a network is an ISN in the case in which there are multiple E cell types. **In a network with multiple E cell subtypes, the network is an ISN if the firing rates of and inhibition received by all E cell types all change in the same direction in response to an input only to I cells. This in turn will be true if all E cell types and all I cell types that connect to E cells all change their steady-state firing rates in the same direction**.

## Discussion

We have established conditions for individual units to have paradoxical steady-state rate responses to stimulation, that is, a steady-state rate response in the direction opposite to that initially driven by the stimulation. Surprisingly, this depends not simply on whether the circuit without that unit is unstable, but on whether it has an odd number of unstable modes. Similarly, a subset of units has an even or odd number of modes with paradoxical responses if the circuit without those units has an even or odd number of unstable modes, respectively.

We have also established that any unstable mode of a subset of units, in response to stimulation of the remaining units, will receive steady-state input from those remaining units that is paradoxical (driving the mode in a direction opposite to its steady-state change in rate), which compensates for an excess of self-input. In particular, any unstable mode of a subset of excitatory cells will, in response to stimulation of inhibitory cells, show a paradoxical change in the inhibition it receives, *i.e.* a change in the same direction as the change in the mode’s amplitude. This means that a network can be shown to be an ISN if the rates of all of the excitatory units, and either the inhibition they each receive or the rates of all of the inhibitory units that project to E cells, all change in the same direction in response to stimulation of a set of inhibitory cells.

We hope that this will clarify the interpretation of future experiments. At the same time, this does not address the effects of heterogeneities in the network or in optogenetic perturbations: *i.e.*, connections among neurons of a given cell type will not all have the same strength, nor will connections between neurons of two given cell types; and optogenetic perturbations may not activate all cells of the targeted type(s), and will stimulate those cells that are acted upon with varying strengths. Thus, a fuller understanding of experiments requires understanding perturbations in high-dimensional models, in which there are many neurons of each cell type with heterogeneous connectivity. We address this in Palmigiano et al. (2020).

## Appendix Interpretation of Eqs. 10

We consider Eq. 10, in which we have changed the basis of the E cells to the basis of eigenvectors of the E→E submatrix of (1 − **W**), or equivalently of **W**.

Let the right and left eigenvectors of the E→E submatrix of **W** be written **e**^*α*^ and **l**^*α*^, respectively, *α* = 1*, …, N*_E_; these are *N*_E_-dimensional vectors. The **W** matrix in this new basis is **E**^−1^**WE**, where **E** is the matrix whose columns are the full N-dimensional basis vectors. The upper left *N*_E_ × *N*_E_ submatrix of 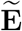 – call it **E** - – is the matrix whose columns are the right eigenvectors **e**^*α*^ of the E→E submatrix of **W**; the lower right *N*_I_ × *N*_I_ submatrix of **E** is the *N*_I_-dimensional identity matrix; and the remaining entries of **E** are zero. Similarly, the upper left *N*_E_ × *N*_E_ submatrix of **E**^−1^ is 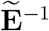, which is the matrix whose rows are the left eigenvectors of the E→E submatrix of **W** written as row vectors, **l**^*α†*^ (where **v**^*†*^ is the complex-conjugate transpose of the column vector **v**); the lower right *N*_I_ × *N*_I_ submatrix of **E**^−1^ is the *N*_I_-dimensional identity matrix; and the remaining entries of **E**^−1^ are zero.

The weight from an I cell *k* to an E mode *α* – the *αk* element of **E**^−1^**WE**, *i.e.* of **W** in the new coordinate system – is then 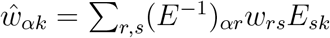. For *k* ≥ *N*_E_ + 1, *E*_*sk*_ = *δ*_*sk*_, so the sum collapses to 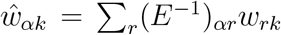 Since 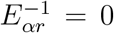 for *α* ≤ *N*_E_ and *r* > *N*_E_, this collapses further to 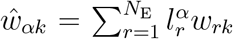 where 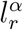 is the *r*^*th*^ element of **l**^*α†*^. Defining **w**^*k*^ as the vector of connection strengths from inhibitory cell *k* to the excitatory cells, with *r*^*th*^ component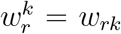, this is 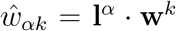. In general, the component of any vector **v** in the direction of eigenvector **e**^*α*^ is given by **l**^*α*^ · **v**. Thus the connection 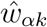 from inhibitory cell *k* to the *α*^*th*^ excitatory mode is precisely the component of **w**^*k*^ in the **e**^*α*^ direction.

The right side of Eq. 10 can then be understood as follows. Define the net inhibition received by the *i*^*th*^ E-cell as 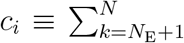, and the *N*_E_-dimensional vector **c** as the vector with *i*^*th*^ element *c*_*i*_. Then the right side of Eq. 10 is

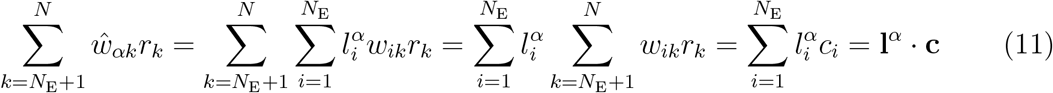

That is, the right side of Eq. 10 is just the component in the direction **e**^*α*^ of the vector **c** of net inhibitory input to excitatory cells.

## Acknowledgements

A.P. acknowledges the support of the Swartz Foundation Fellowship for Theory in Neuroscience 2019-4. This work was supported by funding from NIH 5U19NS107613, U01-NS108683, and R01-EY029999, from NSF NeuroNex 1707398, and from the Gatsby Foundation GAT3708.

## Footnotes

1 We refer to real eigenvalues, because any unstable complex eigenvectors (those with eigenvalues with negative real part) come in complex conjugate pairs, *i.e.* there is an even number of them. The contribution of such a pair to the determinant – the product of the two complex conjugate eigenvalues – is positive. We need not consider these modes; it is the number of modes with real negative eigenvalues that determines whether the determinants are positive or negative. The sense in which such complex conjugate pairs of modes have “paradoxical” response is as follows. Consider two complex conjugate eigenvectors, **e** and **e**^∗^, where the asterisk indicates complex conjugate, with eigenvalues *λ* and *λ*^∗^ respectively. A real stimulus is represented by *c* **e**+ *c*^∗^**e**^∗^ = 2(*c*_*R*_**e**_*R*_ *c*_*I*_ **e**_*I*_), where *c* is a complex number determining the stimulus strength and the subscripts *R* and *I* refer to the real and imaginary part. The response is then 2 (*λ*_*R*_(*c*_*R*_**e**_*R*_ − *c*_*I*_ **e**_*I*_) + *λ*_*I*_ (*c*_*R*_**e**_*I*_ + *c*_*I*_ **e**_*R*_)). If *λ*_*R*_ < 0, this produces a paradoxical mode in the sense that the input 2(*c*_*R*_**e**_*R*_ − *c*_*I*_ **e**_*I*_) produces an output that is a negative multiple of itself, plus a multiple of an additional pattern (*c*_*R*_**e**_*I*_ + *c*_*I*_ **e**_*R*_).

2 See footnote 1 for consideration of complex eigenvalues.

3 It seems unlikely in cerebral cortex that any subtype of excitatory cell does not project to itself or to any of the other excitatory subtypes in the local circuit. However, if there are zeros in the excitatory submatrix, the Perron-Frobenius theorem allows statements to be made that are analogous to those for an all-positive submatrix, although the conclusions will be less air-tight.

## References

Adesnik, H. (2017). Synaptic Mechanisms of Feature Coding in the Visual Cortex of Awake Mice. Neuron, 95:1147–1159.

Garcia Del Molino, L. C., Yang, G. R., Mejias, J. F., and Wang, X. J. (2017). Response reversal during top-down modulation in cortical circuits with multiple interneuron types. Biorxiv, doi: http://dx.doi.org/10.1101/124669.

Karnani, M. M., Jackson, J., Ayzenshtat, I., Tucciarone, J., Manoocheri, K., Snider, W. G., and Yuste, R. (2016). Cooperative Subnetworks of Molecularly Similar Interneurons in Mouse Neocortex. Neuron, 90:86–100.

Kato, H. K., Asinof, S. K., and Isaacson, J. S. (2017). Network-Level Control of Frequency Tuning in Auditory Cortex. Neuron, 95(2):412–423.

Li, N., Chen, S., Guo, Z. V., Chen, H., Huo, Y., Inagaki, H., Davis, C., Hansel, D., Guo, C., and Svoboda, K. (2019). Spatiotemporal limits of optogenetic manipulations in cortical circuits. biorxiv, doi: https://doi.org/10.1101/642215.

Litwin-Kumar, A., Rosenbaum, R., and Doiron, B. (2016). Inhibitory stabilization and visual coding in cortical circuits with multiple interneuron subtypes. J. Neurophysiol., 115:1399–1409.

Mahrach, A., Chen, G., Li, N., van Vreeswijk, C., and Hansel, D. (2019). Mechanisms underlying the response of mouse cortical networks to optogenetic manipulation. arxiv:1907.00816 [q-bio.NC].

Ozeki, H., Finn, I. M., Schaffer, E. S., Miller, K. D., and Ferster, D. (2009). Inhibitory stabilization of the cortical network underlies visual surround suppression. Neuron, 62:578–592.

Palmigiano, A., Fumarola, F., Mossing, D., Kraynyukova, N., Adesnik, H., and Miller, K. D. (2020). Structure and variability of optogenetic responses in cortical circuits with multiple cell-types. bioRxiv, (to be submitted).

Pfeffer, C. K., Xue, M., He, M., Huang, Z. J., and Scanziani, M. (2013). Inhibition of inhibition in visual cortex: the logic of connections between molecularly distinct interneurons. Nat. Neurosci., 16:1068–1076.

Rubin, D. B., Van Hooser, S. D., and Miller, K. D. (2015). The stabilized supralinear network: A unifying circuit motif underlying multi-input integration in sensory cortex. Neuron, 85:402–417.

Sanzeni, A., Akitake, B., Goldbach, H. C., Leedy, C. E., Brunel, N., and Histed, M. H. (2020). Inhibition stabilization is a widespread property of cortical networks. Elife, 9.

Tsodyks, M. V., Skaggs, W. E., and Sejnowski, T. J.and McNaughton, B. L. (1997). Paradoxical effects of external modulation of inhibitory interneurons. J. Neurosci., 17:4382–4388.

